# Large-scale meta-analysis of over one million individuals reveals the genetic architecture of 127 complex traits in East Asian populations

**DOI:** 10.64898/2026.06.18.730290

**Authors:** Jinyeon Jo, Seik-Soon Khor, Shih-Kai Chu, Yunmi Ji, Kazuko Ueno, Akira Ono, Chia-Wei Chen, Ahra Do, Hyein Han, Yousuke Kawai, Nam-Eun Kim, Chun-houh Chen, Katsushi Tokunaga, Sungho Won, Hsin-Chou Yang

## Abstract

Genome-wide association studies (GWASs) have disproportionately focused on European (EUR) populations, limiting the characterization of genetic architecture in other ancestries. To address this imbalance, we integrated large-scale biobanks from Japan, Korea, Taiwan, and China to perform the largest phenome-wide meta-analysis to date in East Asian (EAS) populations, encompassing over one million individuals across 127 complex traits.

We identified 8,010 previously unreported associations and observed substantial genetic sharing across EAS subpopulations, while also detecting cohort-specific heterogeneity within the broader EAS context. Transethnic analyses revealed moderate genetic correlations between EAS and EUR populations, indicating both shared and ancestry-specific components of disease risk. Pleiotropy analyses highlighted prominent signals within the HLA region, supported by protein–protein interaction connectivity and immune-related pathway enrichment. Decomposition of genome-wide association matrices further uncovered structured cross-trait architectures, revealing a predominantly shared polygenic backbone driven by metabolic, biochemical, and anthropometric traits, together with two discrete latent components enriched for immune-related processes.

Together, our findings refine the genetic architecture of complex traits in East Asian populations at unprecedented scale and clarify the balance between shared and population-specific determinants of human diseases.

## Introduction

The Human Genome Project, initiated in 1990 and completed in 2003, established the first reference of the human genome with the goal of comprehensively decoding human genetic information. This foundational effort was followed by large-scale initiatives such as the international HapMap Project, the 1000 Genome (1000G) project and the UK Biobank (UKB), collectively ushering in the era of genome-wide association studies (GWASs). Since the first landmark GWAS identifying susceptibility loci for age-related macular degeneration in 2005 [1], the number of reported associations has expanded rapidly. By September 2016, 24,218 GWAS associations from 2,518 publications had been catalogued [2], and by 2020, over 55,000 loci across nearly 5,000 traits have been reported [3].

Despite this remarkable progress, the majority of large-scale GWASs have been conducted in populations of European (EUR) ancestry, resulting in substantial disparities in representation. As of September 2025, 28.29% of GWASs at the study level and 12.23% of participants at the individual level involved non-EUR populations [4]. This imbalance has constrained the identification of both population-general and population-specific associations and has reduced the predictive accuracy of genetic risk models in non-EUR populations. It has also reinforced a EUR-centric research infrastructure, including the widespread use of EUR-derived summary statistics in polygenic risk modeling under the assumption of moderately shared genetic architecture across ancestries [5, 6], despite improved predictive performance when using meta-analyzed summary statistics derived from target populations such as East Asians (EAS) [7]. Collectively, this disparity not only limits the generalizability of genetic discoveries but also reduces the transferability of findings, including polygenic risk scores across ancestries.

To address this gap, recent efforts have focused on establishing large-scale East Asian cohorts, now representing the second-largest ancestry group in GWAS research. These efforts include Biobank Japan (BBJ) [8], Tohoku Medical Megabank Organization (ToMMo) [9], Korean Cancer Prevention Study-Ⅱ (KCPS2) [10], Korean Genome and Epidemiology Study (KoGES) [11], Taiwan Biobank (TWB) [12] and Taiwan Precision Medicine Initiative (TPMI) [13], and China Kadoorie Biobank (CKB) [14]. While each biobank has contributed valuable insights into EAS subpopulation-specific genetic architecture, most cohorts, except TPMI, remain smaller in scale compared to major EUR resources such as the UKB (approximately 94% of ∼500,000 participants of EUR ancestry) and FinnGen (over 412,000 Finnish participants) [15]. Although prior studies have conducted EAS-specific GWASs and meta-analyses [16–19], restricted access to individual level data and limited availability of comprehensive summary statistics have hindered systematic cross-cohort and cross-trait investigation in EAS populations.

In the present study, in collaboration of Japan, Korea and Taiwan, we performed large-scale meta-analyses of 127 traits across 1,035,254 individuals of EAS ancestry, constructing one of the largest multi-trait genetic analyses in EAS populations to date. By integrating PheWAS-based meta-analyses with downstream analyses of heritability, genetic correlation, gene-based associations, pleiotropy, and latent component (LC) decomposition, we sought to characterize the shared and population-specific genetic architecture of complex traits in EAS populations. These efforts aim to expand genomic resources beyond EUR populations and contribute to ancestry-informed precision medicine.

## Materials and methods

### Study populations

To perform large-scale meta-analyses in EAS populations, we assembled data from five prospective cohorts encompassing four major EAS subpopulations: Japanese (BBJ and ToMMo), Korean (KCPS2, KoGES, and multiple hospital-based cohorts), Taiwanese (Taiwan Biobank; TWB and Taiwan Precision Medicine Initiative; TPMI), and Chinese (CKB). In addition, we included the UKB as a representative EUR cohort and incorporated additional EUR populations from the GWASs catalogue where applicable.

For the Japanese population, we included two Japanese biobank datasets from BBJ (N = 199,982), and ToMMo (N = 87,865) [20]. BBJ is a patient-based cohort launched in 2003 with support from the Japanese Ministry of Education, Culture, Sports, Science and Technology and managed by the Institute of Medical Science at the University of Tokyo and the RIKEN Center for Integrative Medical Sciences. To date, BBJ enrolled 270,000 patients (53.1% male; mean age, 62.7 years for men and 61.5 years for women) with at least one of 51 target diseases from 12 cooperating medical institutions across Japan. ToMMo is a population-based cohort established as a part of the Tohoku Medical Megabank (TMM) project in 2013 by Tohoku University and Iwate Medical University, recruiting residents aged over 20 years from the Miyagi and Iwate prefectures. In addition to its population-based design, ToMMo was established to investigate the multidimensional health effects of the Great East Japan Earthquake and subsequent tsunami, incorporating genetic, environmental, and longitudinal health data from affected regions.

For the Korean population, we integrated two population-based cohorts; KCPS2 (N = 156,701) and KoGES (N = 217,715), and seven hospital-based datasets; the Gene-Environment of Interaction and Phenotype cohort (GENIE; N = 10,300), a subcohort of the Health and Prevention Enhancement study at Seoul National University Hospital Gangnam [21]; the Yonsei University Severance Hospital health checkup cohort (YSUH; N = 6,679); the Veterans Health Service Medical Center medical checkup cohort (VHSMC; N=2,598), gastric cancer (SNUHGC; N =1,216) and lung cancer (SNUHLC; N = 288) cohorts from Seoul National University Hospital; the Kangwon National University Hospital chronic obstructive pulmonary disease cohort (KNUHCOPD; N = 2,115), and the Asan Medical Center Asthma cohort (AMCA, N = 2,046). KCPS2 was initiated in 2004 with the support from the Seoul Metropolitan Government and recruited participants (60.5% male; mean age, 41.7 years), nearly 90% of whom resided in the metropolitan area (Seoul and Gyeonggi Province), through 18 health promotion centers nationwide. KoGES is a government-led population-based cohort initiated in 2001 through collaborations among the Ministry of Health and Welfare, the National Institute of Health, and the Korea Centers for Disease Control and Prevention. It comprises three subcohorts: the Korean Association Resource (N = 10,030; 47.4% male; mean age, 52.3 years), the Cardiovascular Disease Association Study (N = 28,338; 34.2% male; mean age, 53.1 years), and the Health Examinee Study (N = 173,357; 38.19% male; mean age, 58.6 years), enrolling individuals aged over 40 years at baseline.

For the Taiwanese population, we included two population-based cohorts; TWB (N ≈ 150,000) and TPMI (N = 565,390). TWB was initiated in 2012 with support from the Taiwanese government and aims to recruit 200,000 individuals aged 20 - 70 years without a history of cancer [22]. To date, TWB has enrolled 108,955 participants (36.2% male; mean age, 49.9 years), predominantly of Han-Chinese ancestry, through more than 30 recruitment centers across Taiwan. TPMI was established in 2019 through a collaboration among Academia Sinica, 16 partner medical centers, and 33 regional hospitals. Eligibility excluded individuals whose peripheral blood–derived DNA may not accurately represent germline genetic variation, resulting in 486,956 participants (44.7% male; mean age, 57.4 years for men and 54.9 years for women) with available genetic data and linked medical records [13].

For the Chinese population, we included the CKB (N = 512,891) [23], a prospective population-based cohort initiated in 2002 through a collaboration between the Kadoorie Charitable Foundation and UK-based research institutions. Participants (41% male) aged 37-74 years (mean age, 52 years), were recruited between 2004 and 2008 from 10 geographically diverse regions (five urban and five rural) across China.

As a representative EUR cohort, we primarily used the UKB (N ≈ 500,000) [24], a population-based prospective study initiated in 2005 with support from the UK Medical Research Council, the Wellcome Trust, and other funding organizations. UKB recruited participants aged 40-69 years (45.6% male; mean age, 53.5 years), predominantly self-reported as White (94%), through 22 assessment centers across the United Kingdom via the UK National Health Service.

All cohorts included in this study are large-scale, well-characterized biobanks designed to investigate the interplay among genetic, environmental, lifestyle, and socioeconomic factors and health-related outcomes. All participants provided written informed consent for the collection of medical data, biospecimens, and physical measurements in accordance with cohort-specific protocols. Longitudinal follow-up was conducted at the biobank level, typically at intervals of two to five years.

### Genotyping, imputation, and quality control procedures

Most SNP array data were generated using custom-designed genotyping arrays. ToMMo (GRCh38; N ≈ 68,000) was genotyped using the customized Japonica array on the Affymetrix platform [25]. KCPS2 (GRCh37; N = 168,505) was genotyped using either the Illumina Global Screening Array v2.0 or a customized Affymetrix Axiom array v1.0 (Korea Biobank Array; KoreanChip v1.0) [26]. KoGES (GRCh37; N = 72,298) and all Korean hospital-based datasets (GRCh37; N = 7,999 for GENIE and cohort-specific sample sizes for the remaining studies) were genotyped using KoreanChip v1.0 or v1.1. TWB participants were genotyped using two array versions: the Thermo Fisher Axiom (GRCh37 or GRCh38, N = 110,926) with two versions of arrays: the Thermo Fisher Axiom Genome-Wide CHB array (TWBv1; GRCh37; N = 27,719) and a customized Thermo Fisher Scientific array (TWBv2; GRCh38; N = 83,207) [27–29], TPMI (GRCh38; N = 486,956) used two genotyping platforms: TPMv1 (equivalent to TWBv2; N = 165,596) and TPMv2 (an updated version of TPMv1; N = 321,360). Specifically, CKB (GRCh37; N = 105,408) was genotyped using customed CKB arrays on the Affymetrix Axiom platform, based on protocols adapted from the UKB array design [30]. UKB (GRCh37, N = 488,377) was genotyped using two customized arrays: the Applied Biosystems UK BiLEVE Axiom Array (UK BiLEVE; N = 49,950) and the Applied Biosystems UK Biobank Axiom Array (UKB Axiom; N = 438,427).

In contrast, BBJ and a subset of KoGES participants were genotyped using conventional genotyping platform: BBJ (GRCh37, N = 178,726) used either the Illumina HumanOmniExpressExome BeadChip or a combination of the Illumina HumanOmniExpress and the HumanExome BeadChip [31]. A subset of KoGES participants (GRCh37; N = 7,752) was genotyped using Affymetrix Genome-Wide Human SNP Array 5.0 or 6.0.

Phasing and imputation procedures were applied to the summary level data from CKB, BBJ, KCPS2, and TPMI. Briefly, phasing was conducted using Eagle v2.3 [32] for BBJ, SHAPEIT4 [33] for KCPS2, SHAPEIT5 [34] for TPMI, and SHAPEIT3 [35] for CKB. Imputation was performed using IMPUTE4 [36] with the 1000G Project Phase 3 EAS reference panel for CKB; Minimac3 [37] with an integrated reference panel combining 1000G Phase 3 and Japanese whole genome sequencing (WGS) data (N = 1,037) for BBJ [38]; IMPUTE5 with 1000G Phase 3 reference panel for KCPS2; and IMPUTE5 with TWB WGS data (N = 1,498) for TPMI. Detailed quality control (QC) procedures for these datasets are described in the original publications [13, 17, 30, 31]. Publicly available GWAS summary statistics for CKB [23, 39] and BBJ [16] were obtained, and additional downstream QC was applied, excluding single nucleotide polymorphisms (SNPs) with minor allele frequency (MAF) < 0.005. After QC, the final datasets comprised 100,453 Chinese individuals with 8,788,603 SNPs (GRCh37) for quantitative traits, 77,176 Chinese with 8,731,863 SNPs (GRCh37) for categorical traits, 179,000 Japanese individuals with 8,357,933 SNPs (GRCh37), 153,950 Korean individuals with 6,799,813 SNPs (GRCh37), and 342,111 Taiwanese individuals with 8,046,805 SNPs (GRCh38).

For EAS datasets with individual-level genotype (ToMMo, KoGES, Korean hospital-based cohorts, and TWB), biobank-specific processing pipelines were applied using PLINK [40, 41]. For ToMMo, SNPs were excluded based on missing call rate > 0.05, MAF < 0.05, or Hardy-Weinberg Equilibrium (HWE) P < 10^-6^. Individuals were removed for missing call rate > 0.05, sex inconsistencies, excessive homozygosity (> 5 standard deviation (SD) from the mean), or cryptic relatedness (kinship coefficient > 0.0884 [42, 43]). Principal component analysis using the 1000G Phase 3 EAS reference identified two distinct genetic clusters (ToMMo1 and ToMMo2), which were processed separately. Phasing and imputation were performed within each cluster using Beagle (v2) [44] with the customized reference panel combined reference panel of BBJ WGS (N=3,256) and 1000G Phase 3 EAS populations [45]. Post-imputation filtering retained variants with imputation quality R^2^≥ 0.8, yielding final datasets of 42,252 individuals with 6,720,479 SNPs for ToMMo1 and 3,445 individuals with 4,961,913 SNPs for ToMMo2 (GRCh38).

For TWB, we used data phased with SHAPEIT2 [46], imputed with IMPUTE2 [47] using a combined reference panel of TWB WGS (N=1,451) and 1000G Phase 3 EAS populations, followed by TWB-provided post-imputation processing [22]. Additional downstream QC excluded SNPs with missing call rate > 0.05, MAF < 0.005, Bonferroni-corrected HWE P < 1.18 × 10^-8^, or differential missingness between cases and controls for categorical traits. Individuals were excluded for missing call rate > 0.05, sex inconsistency, extreme heterozygosity (> 5 SD from the mean), cryptic relatedness (kinship coefficient > 0.0884), or ancestry outlier relative to the Taiwanese reference cluster based on the first two principal components (PCs). These steps resulted in a final TWB dataset of 108,251 individuals with 4,233,180 SNPs aligned to GRCh38.

KoGES and the seven hospital-based datasets were further stratified by genotyping platform (KoreanChip or Affymetrix) to mitigate batch effects. Data clustering using K-medoid algorithm [48] identified eight clusters, followed by cluster-specific QC and merging. SNP-level QC excluded variants with dish QC < 0.82, missing call rate > 0.03, plate mean call rate < 0.97, plate pass rate < 0.95, duplicated markers, or HWE P < 10^-5^. Individuals were excluded for missing call rate > 0.05, off-target variant heterozygosity < −0.03, extreme homozygosity (< 0.2 or > 0.8), or excessive heterozygosity (> 3 SD from the mean). Platform-specific merged datasets (Affymetrix, VHSMC, KoreanChip, KoreanChip v1.0, and KoreanChip v1.1) underwent additional QC; SNP filtering with a missing call rate > 0.05, excessive heterozygosity (missing call rate, HWE, and MAF), HWE P < 10^-5^, multiallelic or duplicated and individual exclusion with a missing call rate > 0.05, extreme homozygosity (< 0.2 or > 0.8), excessive heterozygosity (> 3 SD from the mean), or duplicated genotypes. Phasing with Eagle v2.4 [49], imputation using the Northeast Asian Reference Database (NARD) imputation server [50] were followed. Post-imputation filtering removed variants with R^2^ < 0.3 or multiallelic sites, resulting in two final merged genotype clusters. For downstream analyses, datasets were stratified by phenotype type and genotyping platform; for SNP extraction, we applied thresholds of missing call rate > 0.05, MAF < 0.05 or 0.005, and HWE P < 10^-5^ or 10^-6^, depending on the datasets. Denoted as Korea Biobank (KRB) in total of 105,792 Koreans, for quantitative traits, we analyzed KRB1_Q_ (87,855 individuals; 4,881,967 SNPs), VHSMC_Q_ (2,433 individuals; 2,943,213 SNPs) from KoreanChip, and KRB2 (7,752 individuals; 2,234,032 SNPs) from Affymetrix. For categorical traits, KRB1_C_ (87,900 individuals; 4,883,024 SNPs) from KoreanChip and KRB2 were used. All Korean datasets were aligned to GRCh37.

For EUR analyses, we used UKB data processed through pre-imputation QC, phasing with SHAPEIT3, imputation with IMPUTE4 with the Haplotype Reference Consortium as the primary reference panel, and UKB-provided post-imputation QC [36]. We restricted analyses to individuals self-identifying as White (responded as White, British, Irish, or any other White background) and applied additional QC excluding variants with MAF < 0.005 and individuals with excessive heterozygosity (> 3 SD from the mean) or ancestry outliers (>3 SD along mean of the first two PCs). The final UKB dataset comprised 392,160 individuals with 4,945,155 SNPs aligned to GRCh37.

### Genome-wide association studies

To construct large-scale genome-wide association studies in EAS populations, we selected traits available in at least two EAS biobanks, as well as nutrition-related traits uniquely available in KRB. This strategy expanded the effective sample size beyond that of previous studies [7]. For ToMMo and KCPS2, analyses were restricted to quantitative traits. The traits were selected based on cross-cohort availability, harmonized phenotype definitions, and sufficient sample size to ensure adequate statistical power for meta-analysis. In total, we analyzed 127 traits, comprising 62 quantitative traits and 65 categorical traits on over one million EAS population.

Categorical disease traits from BBJ, TPMI, and CKB were defined using the ICD10 classification system. In contrast, those from KRB and TWB were not directly ICD-coded, precluding exact cross-biobank matching. For these non-ICD traits, BBJ ICD-based Categorical disease traits from CKB, BBJ, and TPMI were defined using the ICD-based phenotype descriptions were used as the reference framework, and matching was restricted to cases in which the disease name explicitly corresponded to the BBJ ICD trait description to ensure strict phenotype consistency across datasets. To ensure consistency, traits were grouped into higher-level categories based on classification schemes used in prior large-scale EAS PheWAS studies [7, 16, 18]. Quantitative traits were classified into five domains (metabolic, biochemical, anthropometric, reproductive, and nutritional) and categorical traits into six domains (metabolic, digestive, respiratory, neurological, musculoskeletal, and cancer).

Publicly available GWAS summary statistics [16, 17, 19, 30, 39] were generated using linear models (LMs) implemented in PLINK (v1.9/2.0 for CKB), linear mixed models (LMMs) using BOLT-LMM [51] (for BBJ quantitative traits) or SAIGE [52] (v0.37 for BBJ categorical traits, v0.44.5 for KCPS2, and approximately v0.44 for TPMI). For datasets with individual-level genotypes, quantitative traits were inverse-normal transformed to harmonize trait distributions across biobanks and reduce residual population stratification. GWASs were then performed using LMMs in BOLT-LMM (v2.4) for quantitative traits and SAIGE (v1.1.7) for categorical traits, both with construction of genetic relationship matrix with linkage disequilibrium (LD)-independent variants with r2 < 0.2 in each dataset.

De novo GWAS analyses were additionally conducted in UKB only when publicly available EUR summary statistics were unavailable or included fewer participants than our target datasets. All GWAS models included age, sex, and the top ten PCs as covariates.

### Meta-analyses and identification of novel loci

Using GWAS summary statistics harmonized to the GRCh38 genomic assembly, we performed meta-analyses for each EAS subpopulation (Chinese, Japanese, Korean, and Taiwanese) as well as for the combined EAS population using GWAMA (v2.2.2) [53] with a random-effect model modification. To construct EAS PheWASs, we considered two analysis settings to account for difference in study design and population structure: one including all four subpopulations (CJKT; China-Japan-Korea-Taiwan) and another excluding CKB (JKT; Japan-Korea-Taiwan). The exclusion of CKB in the secondary analysis was motivated my its geographically diverse recruitment strategy with reported region-specific variant patterns [54], as well as disease-specific oversampling in the initial CKB array designs from nearly 30% of total genotyped individuals, particularly cardiovascular diseases and COPD [30]. This dual framework allowed us to assess the robustness of meta-analytic findings.

Novel SNP identification was conducted in three steps. First, we defined subpopulation-inclusive SNP sets comprising variants present in at least one large-scale cohort (N > 40,000) within each target EAS subpopulation (7,698,198 SNPs for Chinese, 6,344,259 SNPs for Japanese, 3,350,189 SNPs for Korean, and 3,698,353 SNPs for Taiwanese), according to the CJKT or JKT framework. These sets were then merged to generate an EAS-inclusive SNP list, resulting in 8,583,953 SNPs. Second, we identified LD-independent SNPs using the 1000G Phase3 EAS reference panel by clumping common EAS-inclusive SNPs within 5 Mb windows at an LD threshold of *r^2^* < 0.05 Third, we retrieved all genome-wide significant associations reported in the GWAS Catalog up to December 31, 2025, and matched them to our findings based on rsID or genomic position. Identified SNPs were annotated to genes using standard gene-mapping procedures using ANNOVAR [55].

To determine novelty, we compared our associations against genome-wide significant loci presented in respective publicly available EAS subpopulation GWASs (CKB, BBJ, KoGES, KCPS2, TWB, and TPMI) and listed in the GWAS catalog. The rsIDs and genomic coordinates were lifted over to GRCh38, and gene annotations were harmonized using NCBI gene information from the GWAS catalog. SNPs were considered novel if they did not match any reported associations based on rsID, genomic position (GRCh37/38), or annotated gene symbol.

Unless otherwise specified, all downstream post hoc analyses were based on meta-analysis results derived from the EAS-inclusive SNP set.

### Heritability, genetic correlation, and gene-based analyses

In parallel with the meta-analyses, we estimated SNP-based heritability using linkage disequilibrium score regression (LDSC) [56] for individual EAS datasets, EAS subpopulations, the combined EAS meta-analysis datasets (JKT/CJKT), and EUR population. For categorical traits, liability-scale SNP-based heritability was not estimated because reliable population prevalence estimates were not available across cohorts, and substantial population heterogeneity could introduce bias in liability transformation. Therefore, analyses of categorical traits focused on association discovery rather than liability-scale heritability estimation.

Genetic correlations within EAS (JKT) subpopulations were also assessed using LDSC, and transethnic genetic correlations between EAS (JKT) and EUR populations were assessed using POPCORN [57]. In addition, 35 traits available only in the Korean cohort were excluded from cross-subpopulation genetic correlation analyses, and 33 traits available only in KoGES were excluded from cross-trait genetic correlation analyses across EAS and EUR populations. Consequently, 93 traits were included in cross-subpopulation genetic correlation analyses, and 95 traits were included in cross-trait genetic correlation analyses.

Gene-based association analyses on EAS (JKT) were conducted using MAGMA (v1.10) [58], based on genomic coordinates mapped to GRCh38. Statistical significance was evaluated using Bonferroni-corrected P values. For EAS analyses, SNPs were restricted to the intersection of the predefined EAS-inclusive SNP set and the 1000G Phase 3 EAS reference panel. For EUR analyses, SNPs were restricted to those present in the 1000G Phase 3 EUR reference panel.

### Pleiotropy, network enrichment, and pathway analyses

To characterize shared genetic architectures across 127 traits in the EAS (JKT) population, we leveraged results from meta-analyses and gene-based association analyses. Pleiotropy analyses were conducted at both the variant and gene levels. At the variant level, pleiotropic variants were defined as genome-wide significant variants (P < 5 × 10^-8^) associated with more than 20 traits. At the gene level, pleiotropic genes were defined as genes reaching Bonferroni-corrected significance in gene-based analyses and associated with more than twelve traits. Thresholds at the variant and gene levels were selected to yield comparable numbers of genes between variant-mapped and gene-based pleiotropy analyses. Pleiotropic variants were annotated to genes using ANNOVAR.

Using two sets of pleiotropic genes (variant-mapped genes and gene-based significant genes), we performed protein-protein interaction (PPI) network enrichment and cluster analysis using STRING [59]. For each gene set, genes were submitted to STRING to evaluate functional connectivity based on known and predicted PPIs and to perform PPI enrichment testing. Network clusters were identified using the Markov cluster algorithm (MCL) implemented in STRING with the default inflation parameter (inflation = 3). To evaluate the concordance between clusters identified at the variant and gene levels, we repeated the STRING analysis using the union of pleiotropic genes from both analyses.

Pathway enrichment analyses were performed using clusterProfiler package (v4.14.6) [60] implemented in R (v4.4.1). Significant genes identified from gene-based analyses were tested for enrichment across the three annotation databases: Gene Ontology (GO), Encyclopedia of Genes and Genomes (KEGG), and Reactome. Statistical significance was defined at a false discovery rate (FDR) < 0.05 using the Benjamini–Hochberg procedure.

### Latent component analyses of genetic associations

To assess the underlying genetic architectures represented as LCs in EAS (JKT) populations, we applied truncated singular value decomposition (TSVD) as implemented in the Decomposition of Genetic Associations (DeGAs) framework [61]. As in genetic correlation analyses, we focused on 95 traits existing in more than one EAS subpopulations and applied TSVD on SNP-level association matrices from GWASs.

For TSVD analyses, QC processed 1000G Project Phase 3 reference panel for EAS population was used to construct the reference SNP lists. Variants in 1000G Project were excluded if they were multiallelic, located in the MHC region (chromosome 6: 25,477,797-36,448,354 base pair in GRCh37), had MAF <0.05, HWE P < 10⁻⁵, missing call rate > 0.05, pairwise LD *r^2^* > 0.1. Furthermore, variants with meta-analyses significance P ≥ 0.001 across all traits were also excluded. This filtering yielded 556,921 SNPs for EAS. We collected the Z statistics from meta-analyses results in to a Z score matrix (556,921 SNPs × 94 traits), with substitution of NA values with zero and trait-wise centralization.

The resulting Z-score matrix was decomposed as Z = USV^T^, where U represents variant × LC loadings, S denotes singular values explaining variance, and V corresponds to trait × LC loadings. We set the number of latent components to K = 30 and selected major components that collectively explained more than 70% of the cumulative variance. For each component k, variant-level contribution score 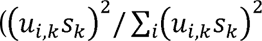 for *i^th^* variant) and trait-level contribution score 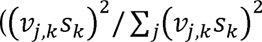, for *j^th^* trait). Gene-level contribution scores were computed by summing variant-level contribution scores of variants mapped to each gene. The top 1% of contributing genes for each component were subsequently subjected to pathway enrichment analyses using the aforementioned methods. By integrating trait contribution and gene contribution scores, we characterized each distinctive LC in terms of its phenotype axis and underlying biological mechanisms. TSVD was performed using the irlba package (v2.3.5.1) [62] in R (v4.1.2).

## Results

### Target sample and trait description

To comprehensively characterize 127 traits across EAS populations spanning eleven categories, we analyzed data from 1,035,254 participants, integrating both summary level and individual-level genotype datasets. Of these, 775,514 participants were derived from CKB, BBJ, KCPS2, and TPMI summary statistics, and 259,740 were from individual-level genotyped datasets including ToMMo, KRB, and TWB. Specifically for KRB, genotype processing was visualized (**SFigure 1**). Cohort characteristics and detailed sample sizes for each analyzed trait are provided in **STable 1-3**.

**Figure 1.**
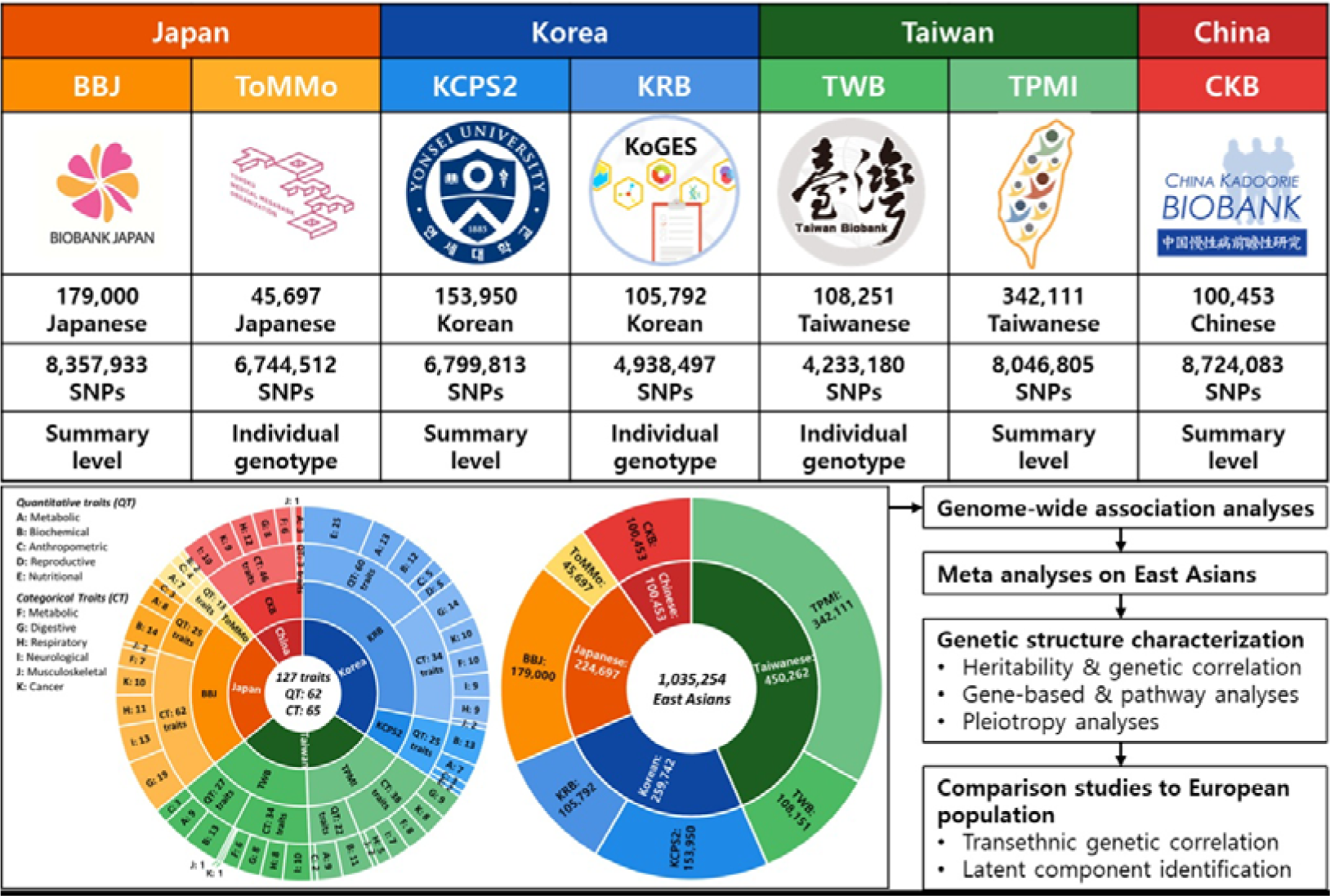
Overview of the study design and analytical workflow. **Note.** BBJ, Biobank Japan; ToMMo, Tohoku Medical Megabank Organization; KCPS2, Korean Cancer Prevention Study-, KRB, Korea Biobank in combination of Korean Genome and Epidemiology Study and multiple hospital-based cohorts; TWB, Taiwan Biobank; TPMI. Taiwan Precision Medicine Initiative; CKB, China Kadoorie Biobank

Among quantitative traits in EAS populations, the largest sample sizes were observed for systolic blood pressure (SBP; N = 837,925), whereas among categorical traits, antihypertensive usage or hypertension (C02; N = 793,831) had the largest sample size. For EUR populations, the largest sample samples were observed for height (Hei; N = 1,597,374) and cirrhosis (Cir; N = 1,580,011), respectively. In total, more than two million individuals were included across EAS and EUR datasets, enabling large-scale EAS and cross-ancestry identification of genetic architecture (**Figure 1**).

### Genome-wide association studies, meta-analyses and identification of novel loci

We first conducted GWASs within each genotyping cluster from individual-level datasets (two clusters from ToMMo, four from KRB, and one from Taiwan), followed by meta-analyses within each EAS subpopulation and across EAS populations (JKT and CJKT).

Across EAS subpopulations, we identified 12,398 genome-wide significant variants across ten traits in Chinese populations, 251,063 variants across 59 traits in Japanese populations, 352,940 variants across 47 traits in Korean populations, and 126,586 variants across 59 traits in Taiwanese populations. After LD clumping, these corresponded to 196, 4,421, 6,397, and 4,048 LD-independent signals in Chinese, Japanese, Korean, and Taiwanese populations, respectively. Most associations across all EAS subpopulations were driven by quantitative traits in category A-C (**SFigure 2** and **STable 4**). Height (Hei; category C) showed the largest number of associations in Japanese, Korean, and Taiwanese populations.

**Figure 2.**
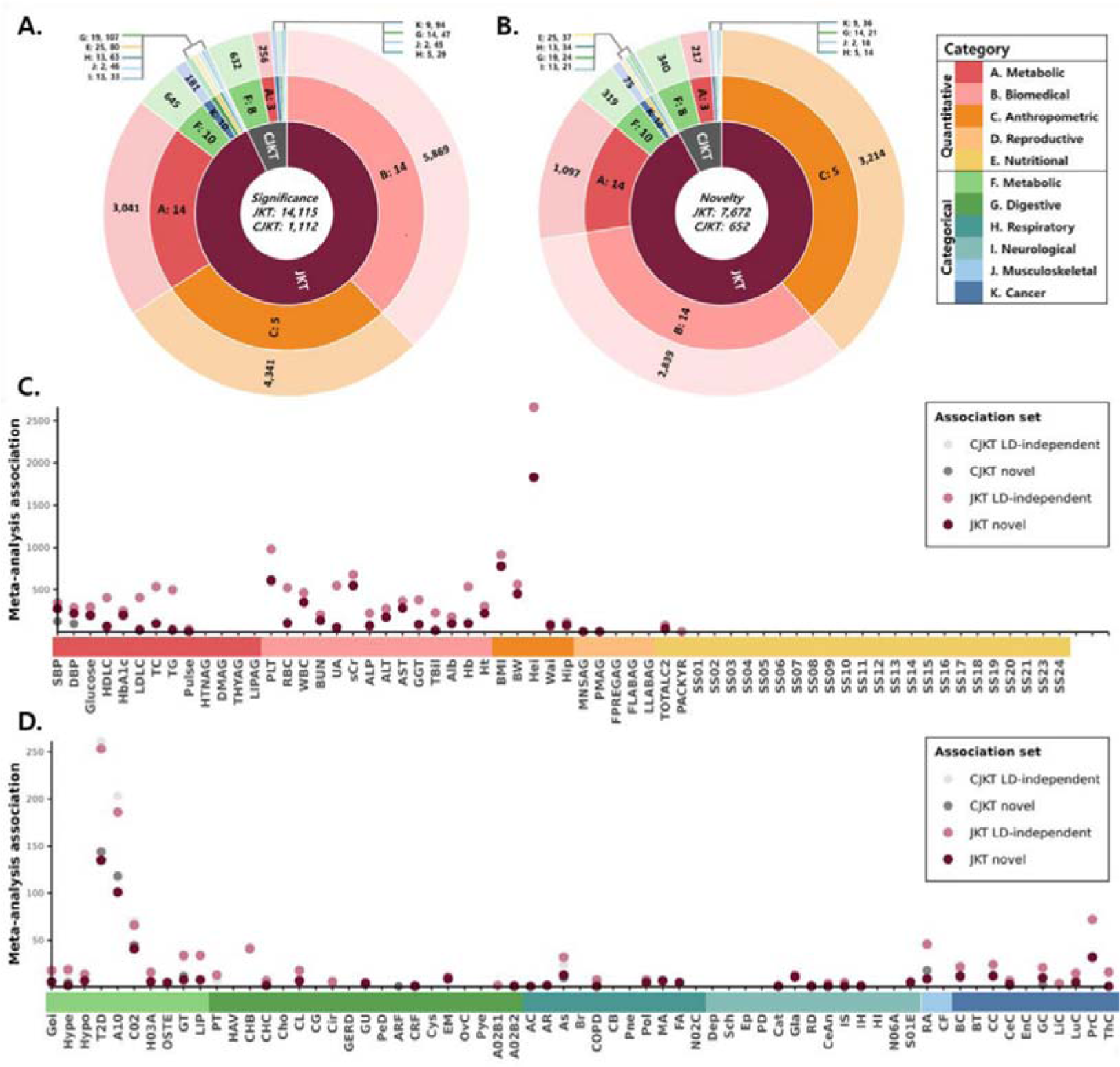
Genome-wide significant and novel associations in EAS meta-analyses. **Note. A and B.** Circular plots summarizing LD-independent genome-wide significant (A) and novel (B) variants identified from two EAS meta-analysis settings (JKT and CJKT). In **A**, LD-independent variants were defined by clumping the overlapping SNP set between the EAS-inclusive SNP list and the 1000 Genomes Project Phase 3 EAS reference panel within 5-Mb windows at an LD threshold of r² < 0.05. The central circle indicates the total number of LD-independent variants per meta-analysis setting; the second ring indicates the EAS meta-analysis framework (JKT or CJKT); the third ring indicates trait categories; and the outer ring indicates the number of LD-independent variants within each category. In **B**, novel variants were defined as those not previously reported in publicly available EAS subpopulation GWASs (CKB, BBJ, KoGES, KCPS2, TWB and TPMI) or in the GWAS Catalog (accessed December 31, 2025), based on rsIDs, genomic position (GRCh37 or GRCh38), or annotated gene symbol. **C and D.** Point plots showing counts of LD-independent genome-wide significant (C) and novel (D) variants from EAS meta-analyses (JKT and CJKT) across quantitative and categorical traits, respectively.

In the EAS-wide meta-analyses, 589,475 variants across 127 traits reached genome-wide significance in JKT, whereas 36,719 variants across 33 traits reached significance in CJKT. Of the CJKT associations, 29,289 variants (79.77%) overlapped with JKT results, corresponding to 45.59% of the 64,246 JKT associations within the same 33 shared traits (**STable 4**). Consistent with the larger effective sample size, JKT yielded a greater number of LD-independent and novel signals than CJKT. Specifically, 14,415 and 1,112 LD-independent variants were identified in JKT and CJKT, respectively, with 551 overlapping signals. For novel associations, 7,672 variants were identified in JKT and 652 variants in CJKT, with 314 overlapping signals. In total, across both EAS meta-analysis settings, we identified 596,905 genome-wide significant associations, including 14,976 LD-independent signals and 8,010 novel associations. Novel loci were predominantly observed for highly polygenic quantitative traits in categories A-C, with additional signals detected in categorical traits within category F (**Figure 2 A-C, STable 5, and STable 6**).

### Heritability, genetic correlation, and gene-based analyses

To characterize genetic architecture within EAS population, we estimated SNP-based heritability and genetic correlations within EAS cohort-level GWASs, subpopulation- and population-level meta-analyses.

#### SNP-based heritability across cohorts

At the dataset level, SNP-based heritability estimates were generally consistent across EAS cohorts for quantitative traits, whereas estimates for categorical traits were uniformly low (**STable 7**). Some smaller datasets (ToMMo2, Bohun, and KoGES) yielded unstable estimates with negative values and large standard error (SE) > 0.2, likely reflecting limited statistical power. Notably, TPMI showed divergent patterns depending on the association model. Estimates derived from LMM (SAIGE) were substantially lower than those obtained from LM (PLINK) and from other EAS cohorts. The PLINK analyses were restricted to unrelated individuals (N = 248,754), whereas the SAIGE analysis included the full sample with extensive relatedness (69.4% of participants having at least one relative closer than third degree) [19]. Correspondingly, SAIGE-based results, in the presence of substantial cryptic relatedness showed deflated test statistics (mean *χ*^2^, λ*_GC_*, and LDSC intercept < 1) [63], while PLINK-based results were mildly inflated (**STable 7**). To minimize inflation and heterogeneity in downstream meta-analyses, we used the LMM-based TPMI results.

#### Heritability in meta-analyses

Across EAS subpopulation and EAS-wide meta-analyses, SNP-based heritability estimates were broadly similar but tended to be modestly lower than those observed in the largest individual cohorts (**STable 8**), consistent with attenuation due to phenotypic and cohort heterogeneity [64]. In the EUR population, estimates were generally higher, likely reflecting larger effective sample sizes and reduced cross-cohort heterogeneity by meta-analyzing a few large-scale cohorts. In both EAS (JKT) and EUR analyses, height (Hei) showed the highest heritability (0.19 in EAS and 0.37 in EUR), whereas categorical traits consistently showed near-zero estimates (**Figure 3A**).

**Figure 3.**
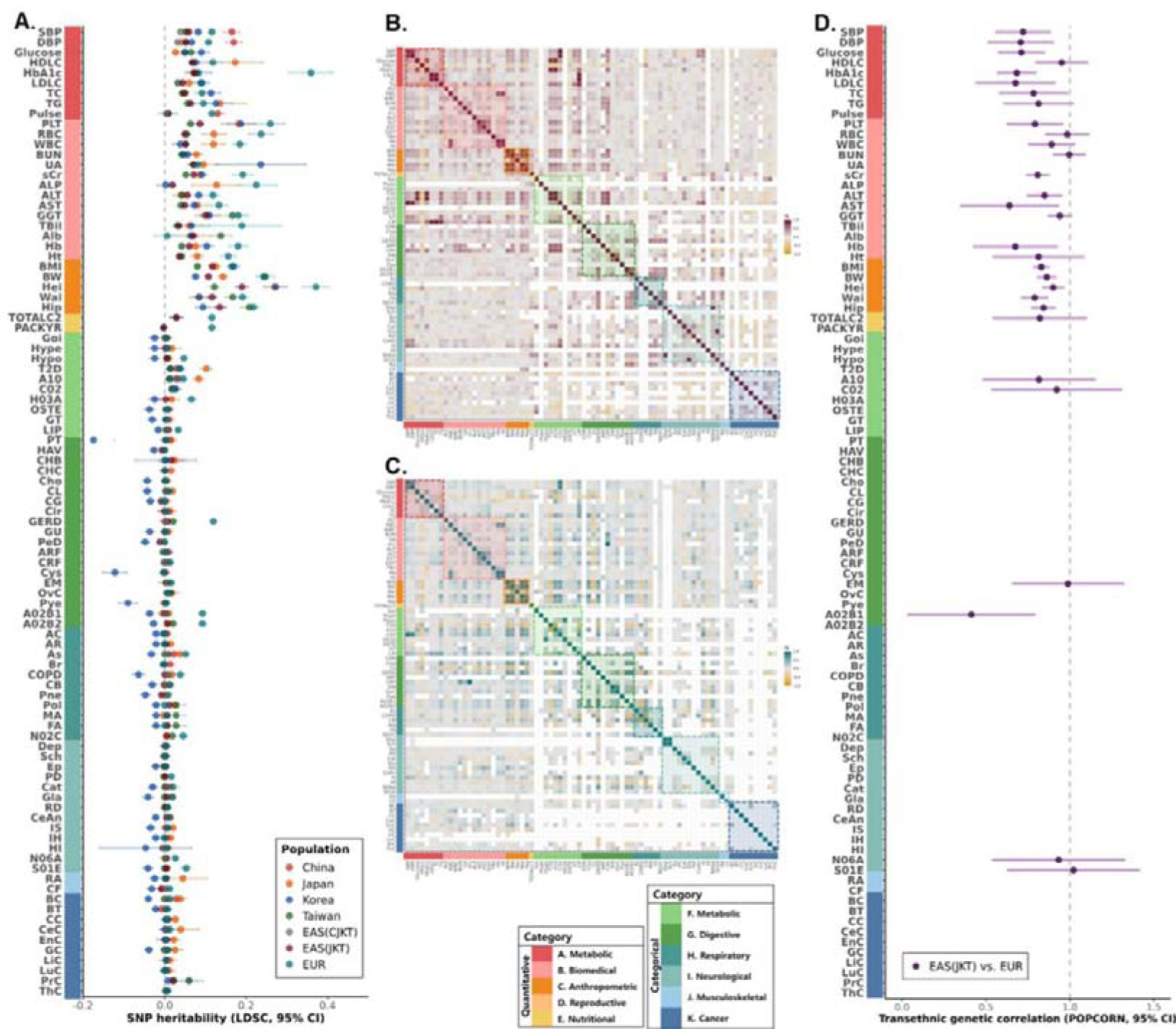
Genetic architecture comparisons across EAS and EUR populations. **Note.** Among the 95 traits existing in more than one EAS subpopulations, SNP heritability and within-ancestry genetic correlation estimates were obtained using LDSC, and transethnic genetic correlations were estimated using POPCORN. Estimates with standard error > 0.2 were excluded from visualization. Error bars indicate 95% confidence intervals. Color bars denote trait categories for both quantitative and categorical traits. **A.** Point plot comparing SNP heritability estimates from EAS and EUR meta-analyses. **B and C.** Genetic correlation matrices of 76 available traits with SE less than 0.2 within EAS (JKT, B) and EUR (C) populations. Strong positive correlations are shown in red (EAS) and blue (EUR), and strong negative correlations in yellow. **D.** Point plot showing transethnic genetic correlations with SE less than 0.2 between EAS (JKT) and EUR populations.

#### Genetic correlations within and across ancestries

Genetic correlations were estimated for 93 traits across EAS subpopulations. Quantitative traits showed consistently high cross-subpopulation correlations, frequently exceeding 0.8 (**STable 9**). In contrast, many categorical traits lacked stable estimates due to low heritability or large SE. Patterns involving the Chinese population (CKB) differed from other EAS subpopulations, which may reflect the geographically diverse recruitment of CKB participants shown in distinct PC patterns [30] and the use of LM–based summary statistics. Given these differences, the JKT meta-analysis was considered the primary representation of shared EAS genetic architecture. Using the EAS-inclusive SNP set, we further evaluated genetic correlations for 76 traits with reliable pairwise estimates (non-negative heritability with SE < 0.2). Similar correlation structures were observed within EAS (JKT) and EUR populations for quantitative traits with SE < 0.2 (**STable 10 and Figure 3B, C**). Transethnic genetic correlations between EAS (JKT) and EUR were also high (approximately 0.8 for traits with SE < 0.2), indicating substantial cross-ancestry sharing of genetic architecture (**STable 11 and Figure 3D**).

#### Gene-based analyses

Gene-based analyses in EAS (JKT) identified 12,899 significant associations across 5,901 genes and 65 traits at Bonferroni-corrected significance. Signals were distributed across all trait categories except D, with 93.6% of associations arising from categories A–C. Height (Hei; category C) alone accounted for 2,451 gene-level associations, consistent with its highly polygenic architecture, along with all of the traits in category C (**STable 12**).

### Pleiotropy, network enrichment, and pathway analyses

We evaluated pleiotropic associations at both variant and gene levels. From MHC inclusive analyses, among 589,475 genome-wide significant associations (379,457 SNPs) from the EAS (JKT) meta-analyses across 76 traits, 39 variants were associated with more than 20 traits and were defined as pleiotropic (**STable 13**). These variants mapped to 38 genes, with one variant unmapped. Associations were predominantly enriched in trait categories A-C, and pleiotropic variants clustered within four genomic regions: *GCKR*/*ZNF512* (chromosome 2), the *HLA* region (chromosome 6), *ABO* (chromosome 9), and *CUX2*/*ATXN2*/*NAA25*/*PTPN11* (chromosome 12) (**Figure 4A**). Notably, 31 of 37 mapped genes localized to or near the *HLA* region. Of the 38 genes, 31 were represented in STRING and were included in protein-protein interaction (PPI) network analyses. These genes formed a significantly enriched interaction network (79 observed edges versus 4 expected; average node degree = 5.27; PPI enrichment P < 1.0 × 10^-16^), indicating strong connectivity. Five interaction clusters were identified (**Figure 4B**), with the *HLA* region cluster connected to the *MICA/MICB* cluster.

**Figure 4.**
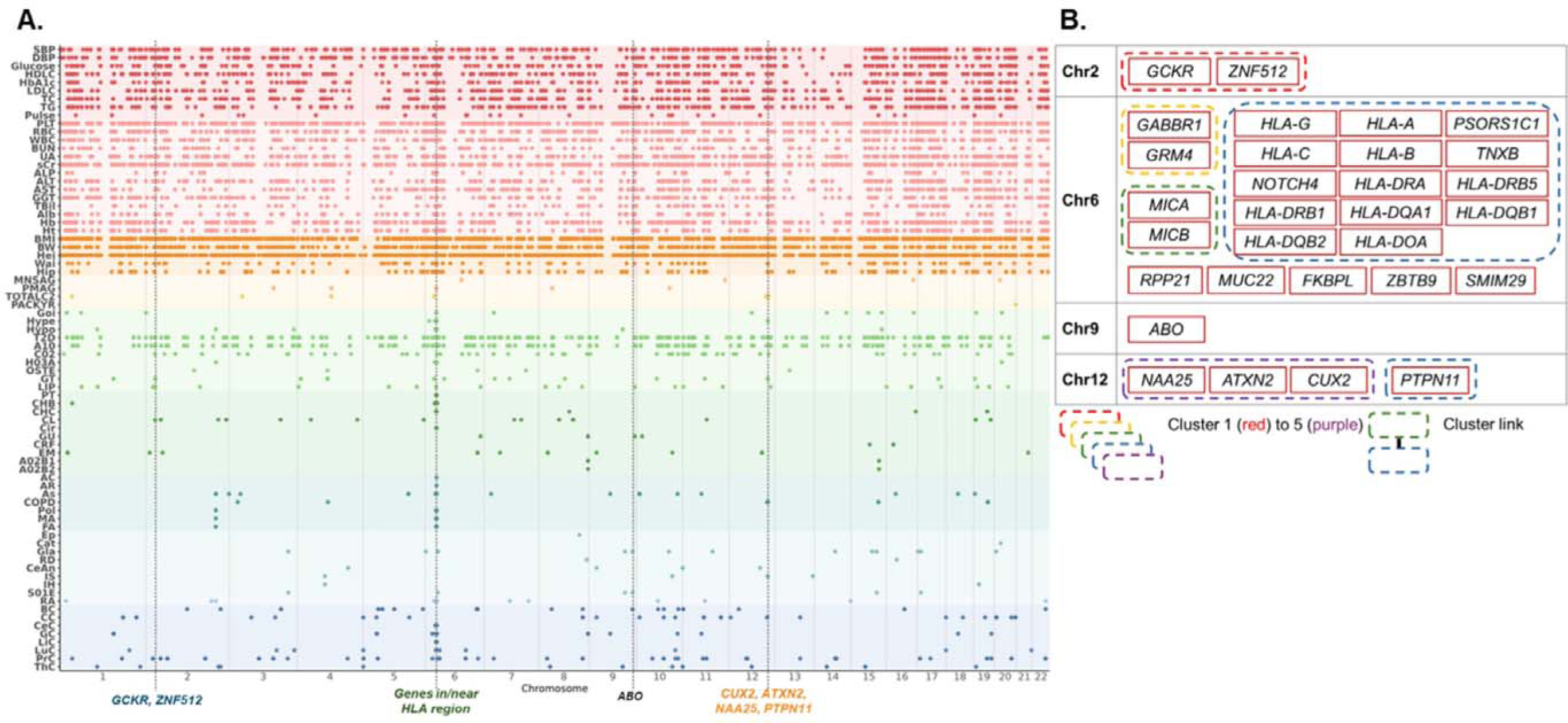
Pleiotropic loci identified in EAS (JKT) meta-analysis and their interaction networks. **Note. A.** Genome-wide significant pleiotropic loci identified from the EAS (JKT) meta-analysis, highlighting four annotated genomic regions. Background colors denote trait categories. **B.** Interaction network of genes matched from pleiotropic loci constructed using STRING. Five interaction clusters were identified among 30 mapped genes from 39 annotated genes.

Gene-based analyses identified 12,899 significant gene-trait associations across 5,901 genes and 65 traits, predominantly in categories A-C. 31 genes were associated with more than twelve traits and were defined as pleiotropic (**STable 14**). These genes spanned eight chromosomes and included all pleiotropic regions identified at the variant level (**SFigure 3A**), with 12 genes overlapping between the two approaches. STRING analyses of these 31 genes also demonstrated significant network enrichment (32 observed edges versus 4 expected; average node degree = 2.06; PPI enrichment P < 1.0 × 10^-16^). Five clusters were identified, two of which overlapped with variant-level clusters, indicating partial concordance between variant-level and gene-level pleiotropy. Notably, *GCKR* (chromosome 2) and *FTO*/*GIPR* (chromosome 16/19) were assigned to distinct clusters but remained functionally connected within the broader interaction network (**SFigure 3B**), suggesting modular organization of pleiotropic effects.

STRING analyses of the union set of pleiotropic genes included 31 genes from each level, with twelve genes overlapping between variant and gene level analyses. Among gene level pleiotropic genes, 29 corresponded to variant level pleiotropic genes (minimum = 8, median = 16, mean = 19.45, maximum = 42 trait associations), whereas 29 variant level pleiotropic genes corresponded to gene level pleiotropic genes (minimum = 1, median = 11, mean = 9.55, maximum = 21 trait associations). In total, 49 unique genes were represented and formed a significantly enriched interaction network (105 observed edges versus 11 expected; average node degree = 4.29; PPI enrichment P < 1.0 × 10^-16^). Eight clusters were identified, with four inter-cluster connections (**SFigure 5**). Most clusters were preserved relative to the variant and gene level analyses, except for cluster 5 (*MUC22*/*TCF19*), which merged variant and gene level signals.

**Figure 5.**
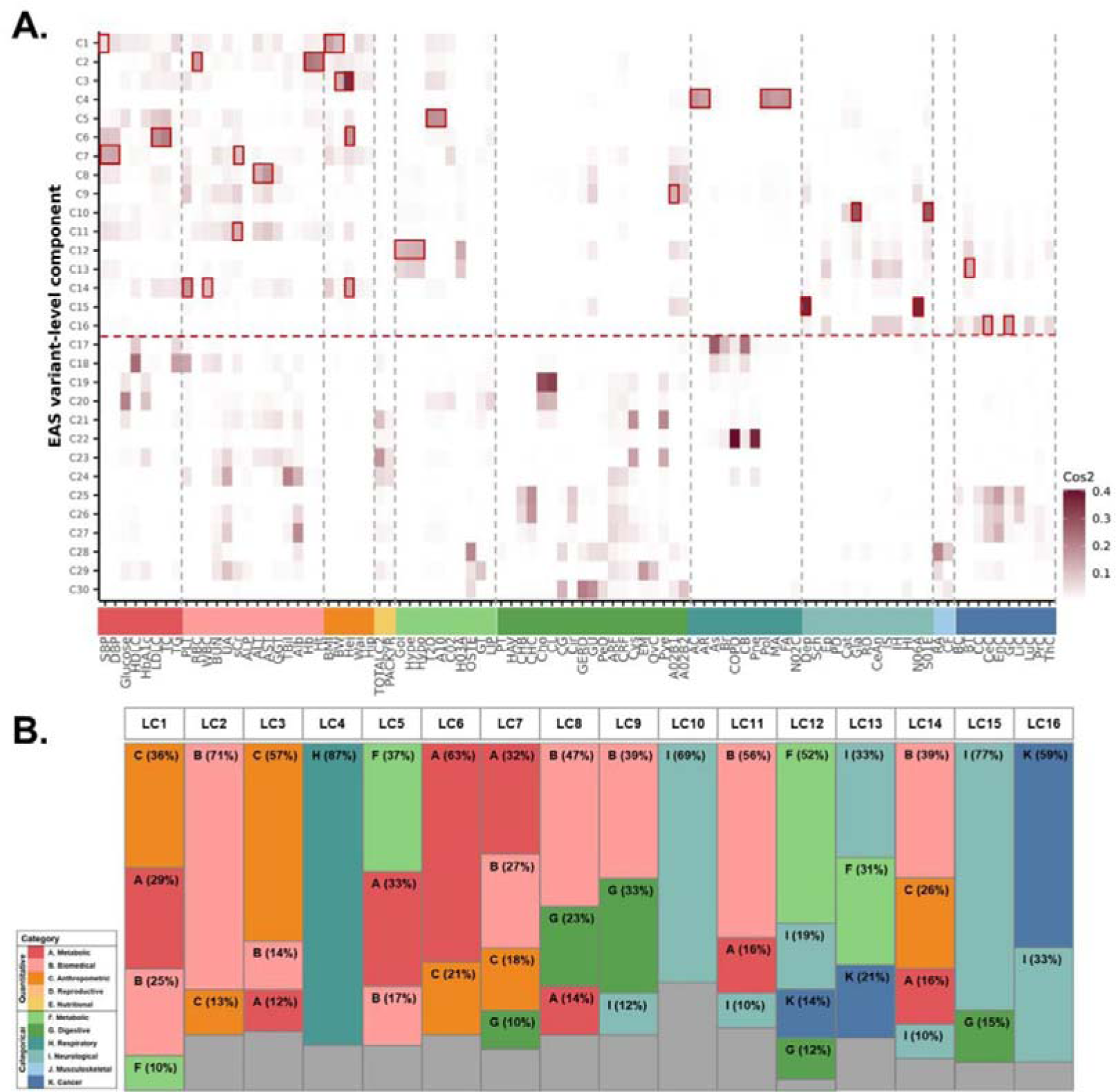
Trait contribution score patterns and category-level contributions across latent components. **Note. A.** Red boxes indicate trait contribution score > 0.1. The red dashed line marks the first 16 latent components, which collectively explain 70% of the total variance. For each latent component, trait contribution scores sum to one. **B.** Category-level contributions are shown for trait categories explaining more than 10% of the total contribution within each latent component. Bars are colored according to the designated category colors, and the gray represents the combined contribution of all remaining categories

Pathway enrichment analyses of 65 traits with significant gene-based associations identified 5,407 enriched pathways (**STable 15**). The largest number of enriched pathways was observed for rheumatoid arthritis (RA; 287 pathways) at the individual-trait level, for category B (biochemical traits; 1,099 pathways) at the trait-category level, and within the GO database (3,989 pathways; 73.77%) among the annotation database. Enrichment was dominated by immune-related pathways, particularly those related to viral infection, antigen processing and presentation, and autoimmune diseases, consistent with the strong contribution of HLA-associated loci.

### Latent component analyses of genetic associations

LCs were selected based on a cumulative explained variance threshold of 70%, resulting in 16 retained components (**SFigure 5**). Trait contribution scores were computed for each LC, with contributions summing to one within each component (**STable 16**). Across the 16 LCs, trait contributions were broadly distributed among categories A-C in most components, reflecting shared polygenic architecture across metabolic, biochemical, and anthropometric traits (**Figure 5A**). In contrast, several components exhibited concentrated trait contributions. LC10 was primarily driven by glaucoma-related traits (Gla: 0.29 and S01E: 0.29), whereas LC15 was dominated by depression-related traits (Dep: 0.37 and N06A: 0.37), indicating disease-specific latent structure. At the category-level contributions were predominantly enriched in category A-C across components. However, specific LCs showed dominance of other categories, including respiratory (H), neurological (I), and cancer-related traits (K), particularly in LC4, LC10, LC15, and LC16 (**Figure 5B**), suggesting modular organization of trait domains within the latent space.

Variant contribution scores aggregated to genes revealed relatively diffuse patterns across components, with several genes contributing broadly across multiple traits (*CSMD1*, 30 traits; *RBFOX1*, 29 traits; *PTPRD*, 26 traits, and *LINGO2*, 24 traits among the top ten contributing genes) (**STable 17**). These widespread contributions nevertheless formed biologically coherent clusters when examined through pathway enrichment analyses (**STable 18**). Notably, components dominated by specific trait categories exhibited distinct functional enrichment patterns. The top 1% contributing genes of LC4, LC10, LC15, and LC16 were significantly enriched in immune regulatory pathways, retinal neurodevelopment processes, cortical excitatory neurotransmission, and oncogenic susceptibility pathways, respectively, providing biological interpretability to disease-focused LCs.

## Discussion

In this study, we constructed large-scale PheWAS meta-analyses of 127 traits across eleven categories in over one million individuals of EAS ancestry. By integrating SNP-based heritability, cross-trait and cross-population genetic correlations, transethnic genetic correlations, MAGMA gene-based association analyses, pleiotropy assessment, pathway enrichment analyses, PPI network analyses, and LC decomposition using the DeGAs framework, we sought to characterize the shared and population-specific genetic architecture of complex traits in EAS populations.

We observed negative genetic correlations between the Chinese (CKB) cohort and other EAS subpopulations for multiple traits (**STable 9**), indicating heterogeneity within the broader EAS population. While EAS subpopulations share substantial ancestral background, important differences arise from study design and internal population structure. Specifically, TWB participants represent a relatively homogenous genetic structure, with over 99% of Han Chinese descent and limited substructure beyond historical migration gradients. In contrast, CKB incorporates participants recruited from ten geographically diverse regions across China, capturing considerable within-China population substructure and region-specific polymorphic variation. Together with disease-specific oversampling in the initial CKB array design, these factors may have contributed to divergence in genetic correlation estimates, might reflecting observed differences in cohort composition and internal heterogeneity rather than fundamental ancestry divergence across EAS populations.

Accordingly, we performed meta-analyses with and without the CKB cohort (CJKT and JKT). Despite the larger sample size, the CJKT analyses yielded fewer significant associations than JKT, with approximately 79.77% overlap. This discrepancy may reflect distinct characteristics of the CKB cohort, including regional stratification across multiple geographic areas and differences in modeling strategies. In TPMI, where substantial cryptic relatedness was present, the use of LMMs resulted in mildly deflated test statistics, whereas LMs applied to unrelated subsets produced estimates comparable to those from other non-Chinese cohorts (mean *χ*^2^, λ*_GC_*, and LDSC intercept; **STable 7**). These observations underscore the importance of cohort-specific structure and modeling strategy in large-scale meta-analyses. Across the two EAS meta-analysis settings, we identified 596,905 genome-wide significant associations, including 8,010 putatively novel associations. SNP-based heritability estimates and cross-population genetic correlations suggested broadly shared genetic architecture among EAS subpopulations (excluding CKB) and between EAS (JKT) and EUR populations (**Figure 3**). Although heritability estimates in EAS were generally lower than in EUR, this pattern may partly reflect attenuation due to meta-analyses across heterogeneous cohorts, as previously reported [64]. Gene-based aggregation using MAGMA identified 5,901 genes across 65 traits, predominantly within metabolic, biochemical, and anthropometric categories, consistent with their highly polygenic architecture.

At the pleiotropic level, both variant- and gene-based approaches identified convergent signals in several genomic regions, including *GCKR*/*ZNF512* (chromosome 2), the *HLA* region (chromosome 6), *ABO* (chromosome 9), and *CUX2* (chromosome 12). Gene-level pleiotropy encompassed a broader set of loci compared to variant-level analyses, reflecting the aggregation of SNP effects. When analyzed separately or jointly in STRING, both pleiotropic gene sets formed significantly enriched PPI networks, indicating non-random functional connectivity. While most clusters were largely preserved across analytical strategies, integration of the two gene sets revealed additional cluster (cluster 4 in **SFigure 5**) and connection (between cluster 4 and cluster 5 in **SFigure 5**), suggesting complementary information captured by meta-analysis and gene-based approaches.

The prominent role of HLA genes within these networks highlights their function as pleiotropic hubs influencing diverse human traits, spanning biochemical, immunological, and disease-related phenotypes [16, 65–68]. However, given the extensive LD structure within the MHC region, such co-occurrence likely reflects both functional immune pathways and shared haplotypic architecture. Connections between clusters may therefore represent a combination of biological interaction and LD-mediated correlation. For example, cluster-level relationships involving metabolic genes between *GCKR* (cluster 1; glucose regulation) [69, 70] and *FTO* (cluster 2; adiposity) [71, 72] / *GIPR* (cluster 2; incretin signaling) [73], as previously reported. Similarly, reported epistasis interactions between *MUC22* and *TCF19* (both in cluster 4, MHC class Ⅰ genes) in type 1 diabetes [74], and residual effects of *TCF19* after adjusting HLA class Ⅱ haplotypes on T1D [75], suggest that non-HLA genes within the MHC region may contribute to disease risk beyond classical antigen presentation loci. In addition, the clusters 3 (HLA-DR- and HLA-DQ-, HLA class Ⅱ genes) and cluster 5 (all four genes, MHC class Ⅲ genes) with connectivity were previously reported STRING analyses on chronic obstructive pulmonary diseases [76]. Variants in *NOTCH4* and *TNXB* (both in cluster 5, MHC class Ⅲ genes) have been associated with regulatory effects on HLA class Ⅱ gene expression in immune-response contexts, supporting the presence of regulatory connectivity across the broader MHC locus [77]. The *MICA* and *MICB* genes (MHC class Ⅰ genes) in cluster 6, although structurally related to classical HLA class Ⅰ molecules, function as stress induced ligands rather than antigen-presenting molecules [78], further illustrating the layered immune architecture embedded within the MHC region. Pathway enrichment analyses revealed that immune-related pathways, particularly viral infection such as HTLV-1 (hsa05166) and Epstein-Barr virus infection (hsa05169), were among the most significantly enriched. This pattern is largely driven by strong pleiotropic effects within the MHC region. While these findings reinforce the centrality of immune-related pathways, disentangling locus-specific functional effects from LD-driven enrichment signals requires further investigation.

To explore higher-order structure across traits, we applied TSVD to derive LCs. To minimize bias from complex LD, the MHC region was excluded from LC analyses. *CSMD1* (chromosome 8) and *RBFOX1* (chromosome 16), contributed broadly across multiple LCs (**STable 17**). *CSMD1*, located outside the MHC region, is implicated in complement-mediated immune modulation and neurodevelopment processes [79], whereas *RBFOX1* functions as a key regulator of alternative splicing in cardiac and neural disorders [80, 81]. Their widespread contribution across components may reflect involvement in fundamental biological processes influencing multiple phenotypic domains. Although many LCs were not readily interpretable in a biologically specific manner, several showed coherent patterns (**Figure 5A/5B** and **STable 18**). LC4 was enriched for airway immune inflammation (e.g., asthma and allergic rhinitis), with gene enrichment: supporting immune activation and T-cell mediated responses. LC10 was characterized by ocular traits such as glaucoma and cataract, with enrichment in synaptic signaling and extracellular matrix pathways. LC15 reflected psychiatric liability, particularly depression, with enrichment in synaptic transmission and glutamatergic signaling pathways. LC16 was enriched for multiple cancer traits and associated pathways related to DNA repair and immune evasion. These findings suggest that TSVD-based decomposition can partially recover biologically meaningful axes of genetic variation across complex traits of the EAS population.

Several limitations should be acknowledged. First, residual heterogeneity across cohorts, stemming from differences in participant collection design, genotyping platforms, imputation reference panels, QC procedures, and regression modeling, may influence meta-analysis results. Second, we primarily relied on LDSC for SNP-based heritability and genetic correlation estimation. Alternative methods for heritability estimation such as S-LDSC [82], LDAK [83], and HDL [84], or local genetic correlation estimation such as rho-HESS [85] and LAVA [86] may provide complementary insights for polygenicity and correlated regions. In addition, robust and widely adopted methods for transethnic local genetic correlation remained limited, restricting fine-scale cross-ancestry comparison. Finally, while immune-related signals were pervasive, particularly within the MHC region, detailed functional dissection of immune and complement pathways with adjustment for LD requires further investigation beyond summary-level analyses. Despite these limitations, this study represents one of the largest PheWAS meta-analyses conducted in EAS populations to date. By integrating multi-layered genomic analyses, we provide a comprehensive overview of shared and population-specific genetic architecture across a broad spectrum of human traits. The identification of 8,010 novel associations underscores the importance of expanding large-scale genetic studies beyond European populations. Together, our findings contribute to the development of EAS-focused genomic resources and provide a foundation for future fine-mapping, functional, transethnic investigations, and EAS-specific precision medicine.

## Data availability

**Summary-level data:** Access to CKB genotype data can be requested through the official website (https://www.ckbiobank.org/) and summary statistics are publicly available at https://pheweb.ckbiobank.org/phenotypes; Access to BBJ genotype data can be requested through the official website (https://biobankjp.org/) and summary statistics are publicly available at https://pheweb.jp/phenotypes; KCPS2 genotype is not publicly available but summary statistics are publicly available at https://zenodo.org/records/15132424; Access to TPMI genotype data can be requested through official website (https://tpmi.ibms.sinica.edu.tw/) and summary statistics are publicly available at https://pheweb.ibms.sinica.edu.tw/. The EUR summary statistics are publicly available in GWAS Catalog (https://www.ebi.ac.uk/gwas/) or Neale lab (https://pheweb.org/UKB-Neale/).

**Individual-level data:** Access to ToMMo genotype data is not publicly available (https://www.megabank.tohoku.ac.jp/) and summary statistics will be uploaded upon publicaion. Access to KoGES genotype data can be requested through official website (https://coda.nih.go.kr/) and the rest of Korean genotype data are not publicly available. Access to TWB genotype data can be requested through official website (https://www.biobank.org.tw/). The summary statistics from Korean data and TWB will not publicly available due to the data policy. Access to UKB genotype data can be requested through official website (https://www.ukbiobank.ac.uk/) and the summary statistics are available upon request. All EAS meta-analyses summary statistics will be uploaded in GWAS Catalog upon publication.

## Ethics statement

The Institute Review Board at Academia Sinica approved our study (AS-IRB01-17049 and AS-IRB01-24070). The Institute Review Board at Seoul National University approved our study (E2303/004-012). The use of ToMMo data has been approved under the grant JIHS-A-003267.

## Acknowledgements

We sincerely appreciate the dedication and support of our project members.

**Taiwan:** We acknowledge research grants from the Academia Sinica (AS-SH-112-01 and AS-FILBD-114-05). We acknowledge the Taiwan Biobank for providing the data and appreciate the invaluable contribution of its participants (TWBR10704-05, TWBR10810-06, TWBR10911-01, and TWBR11406-05). We are grateful to all the participants of the Taiwan Biobank and Taiwan Precision Medicine Initiative Projects.

**Korea:** This work was supported by the National Research Foundation of Korea (NRF) grant funded by the Korea government (MSIT) (RS-2024-00346850). This work was supported by Institute of Information & communications Technology Planning & Evaluation (IITP) grant funded by the Korea government (MSIT) (No.RS-2025-25441324, Development of AI-Enabled, Phenome Data-Driven Precision Health Prediction and Management Solutions for Customized Life Coaching of Healthy Adults). We are grateful to all the participants of the Korea Biobank Projects.

**Japan:** This research was supported by Japan Agency for Medical Research and Development (AMED) under the Genome Research and Innovation Network Platform (GRIFIN) program. This research used data provided by the Tohoku Medical Megabank Organization (ToMMo), Tohoku University. This research was supported by Japan Agency for Medical Research and Development (AMED) under the Genome Research and Innovation Network Platform (GRIFIN) program (17km0405205h0002). We are grateful to all the participants of the Tohoku Medical Megabank Project.

## CRediT author statement

**Jinyeon Jo:** Conceptualization, Methodology, Software, Validation, Formal analysis, Investigation, Data Curation, Writing - Original Draft, Writing - Review & Editing, Visualization, and Project administration. **Seik-Soon Khor:** Conceptualization, Resources, Writing - Review & Editing, Project administration, and Supervision. **Shih-Kai Chu:** Data Curation, Formal analysis, and Writing - Review & Editing. **Yunmi Ji:** Data Curation, Formal analysis, and Writing - Review & Editing. **Kazuko Ueno:** Data Curation, Formal analysis, and Writing - Review & Editing. **Akira Ono:** Data Curation, Formal analysis, and Writing - Review & Editing. **Chia-Wei Chen:** Data Curation and Formal analysis. **Ahra Do:** Data Curation. **Hyein Han:** Data Curation. **Yousuke Kawai:** Conceptualization. **Nam-Eun Kim:** Data Curation. **Chun-houh Chen:** Resources, Writing - Review & Editing, Project administration, and Funding acquisition. **Katsushi Tokunaga:** Resources, Writing - Review & Editing, and Funding acquisition. **Sungho Won:** Conceptualization, Resources, Writing - Review & Editing, Supervision, and Funding acquisition. **Hsin-Chou Yang:** Conceptualization, Resources, Writing - Review & Editing, Supervision, Project administration, and Funding acquisition.

## Supplementary Figures

**Supplementary Figure 1.**
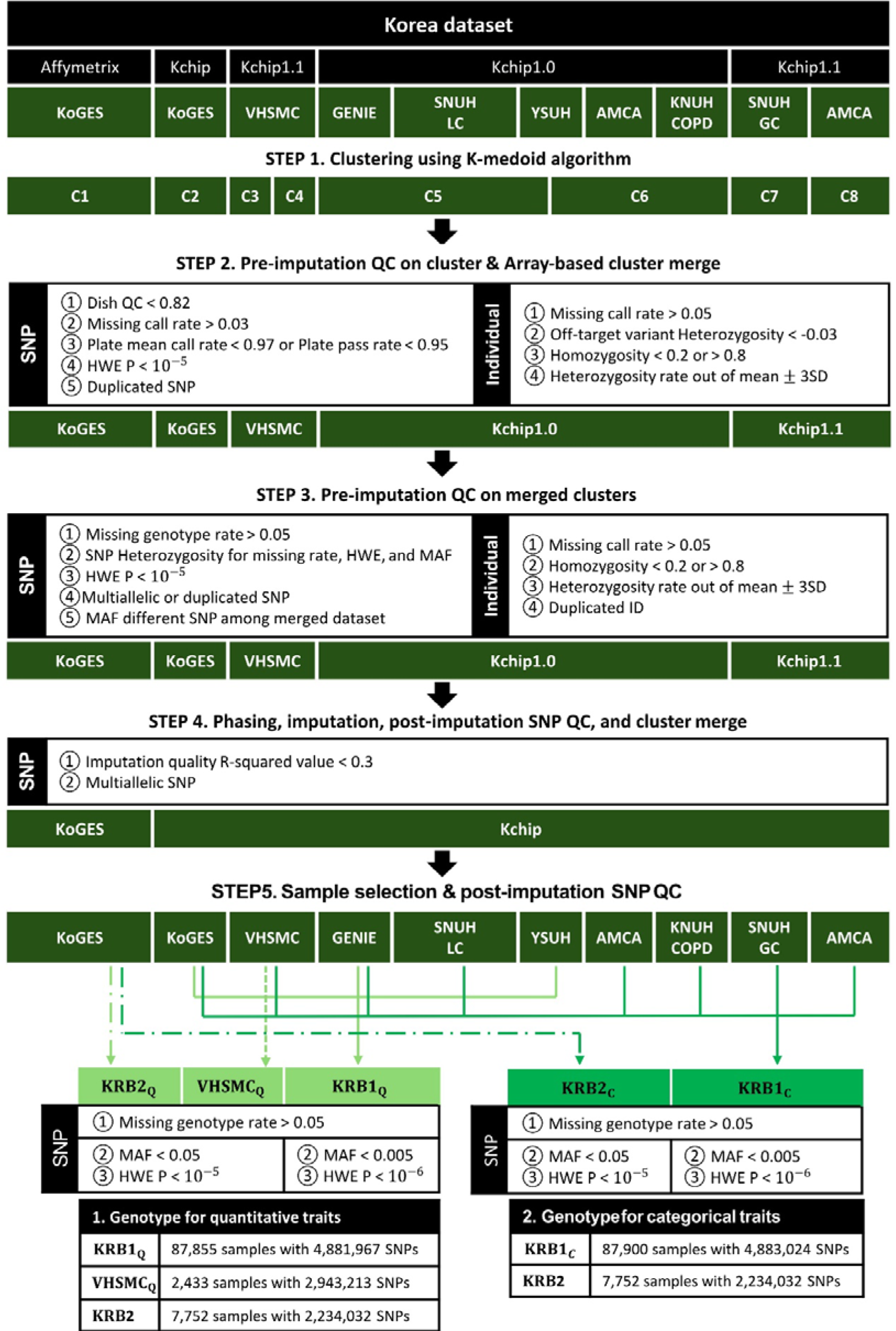
Genotyping and quality control workflow for the Korea Biobank. **Note.** Kchip, Korea Biobank Array (KoeranChip); KoGES, Korean Genome and Epidemiology Study; VHSMC, Veterans Health Service Medical Center health checkup cohort; GENIE, Gene–Environment of Interaction and Phenotype cohort; SNUHLC, Seoul National University Hospital lung cancer cohort; YSUH, Yonsei University Severance Hospital health checkup cohort; AMCA, Asan Medical Center asthma cohort; KNUHCOPD, Kangwon National University Hospital chronic obstructive pulmonary disease cohort; SNUHGC, Seoul National University Hospital gastric cancer cohort; HWE, Hardy–Weinberg equilibrium; MAF, minor allele frequency; KRB, Korea Biobank.

**Supplementary Figure 2.**
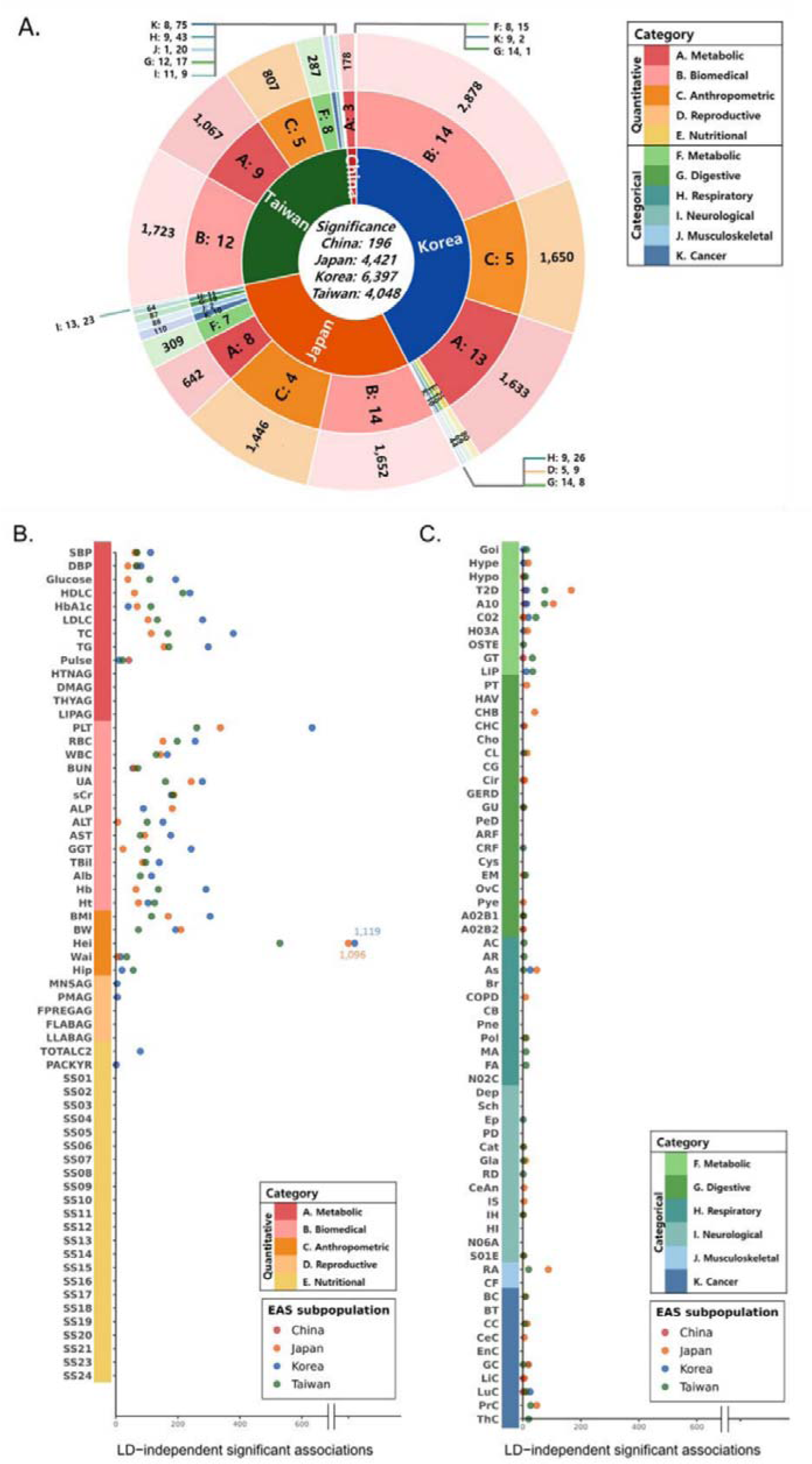
Genome-wide association comparisons from each EAS subpopulation meta-analysis. **Note. A.** Circular plot summarizing LD-independent genome-wide significant variants from each EAS subpopulation meta-analysis. LD-independent variants were defined by clumping the overlapping SNP set between the EAS-inclusive SNP list and the 1000 Genomes Project Phase 3 EAS reference panel within 5 Mb windows at an LD threshold of r² < 0.05. The central circle indicates the total number of LD-independent variants per subpopulation; the second ring indicates EAS countries; the third ring indicates trait categories; and the outer ring indicates the number of LD-independent variants within each trait category. **B and C.** Point plots showing counts of LD-independent genome-wide significant variants across quantitative and categorical traits, respectively.

**Supplementary Figure 3.**
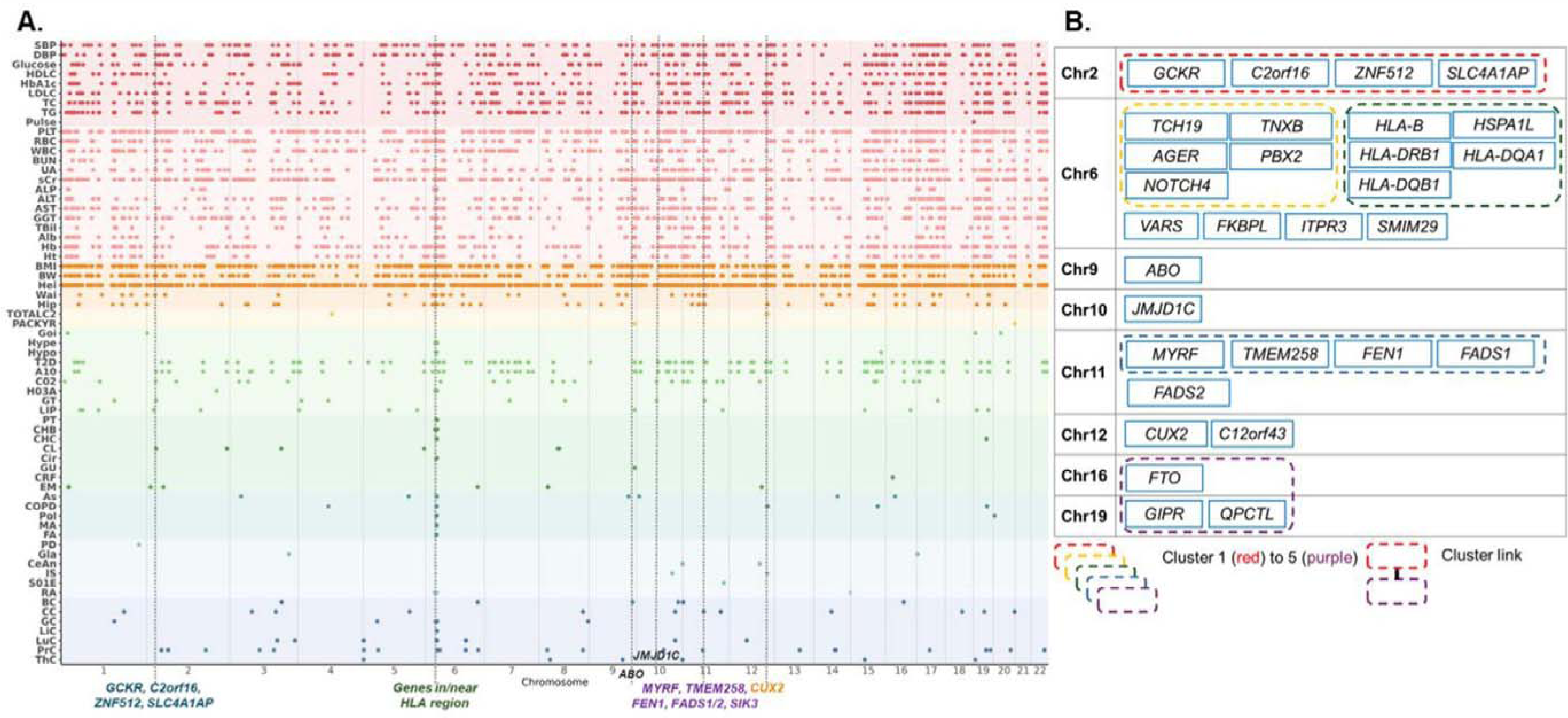
Pleiotropic genes identified in the EAS (JKT) gene-based analysis and their interaction networks. **Note.** *C6orf1* gene was mapped to *SMIM29*. **A.** Bonferroni corrected significant pleiotropic genes identified from the EAS (JKT) gene-based analysis, highlighting six genomic regions. Background colors denote trait categories. **B.** Interaction network of genes constructed using STRING. Five interaction clusters were identified among 31 genes from 31 pleiotropic genes.

**Supplementary Figure 4.**
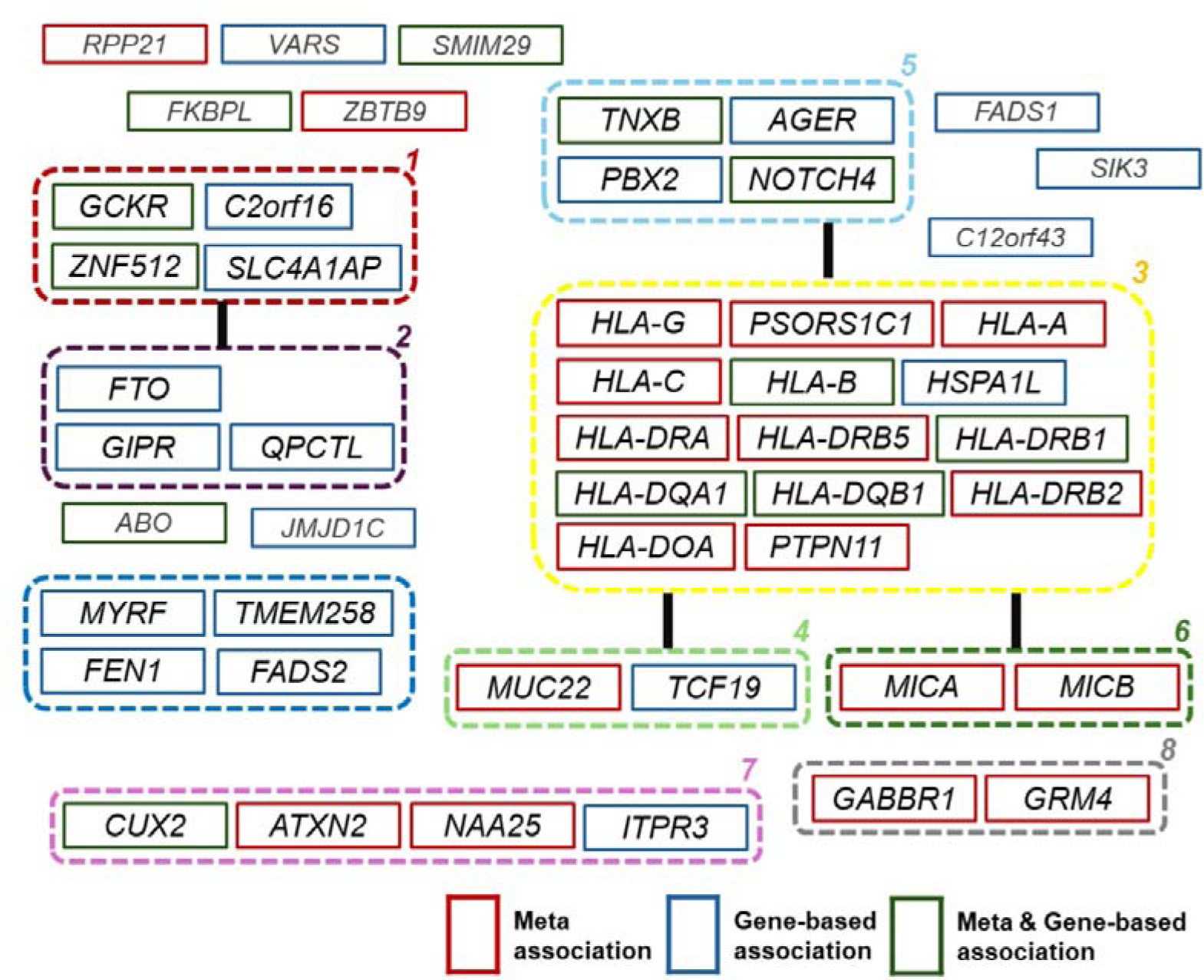
Integrated pleiotropic genes from EAS (JKT) meta- and gene-based analyses and their protein–protein interaction networks.

**Supplementary Figure 5.**
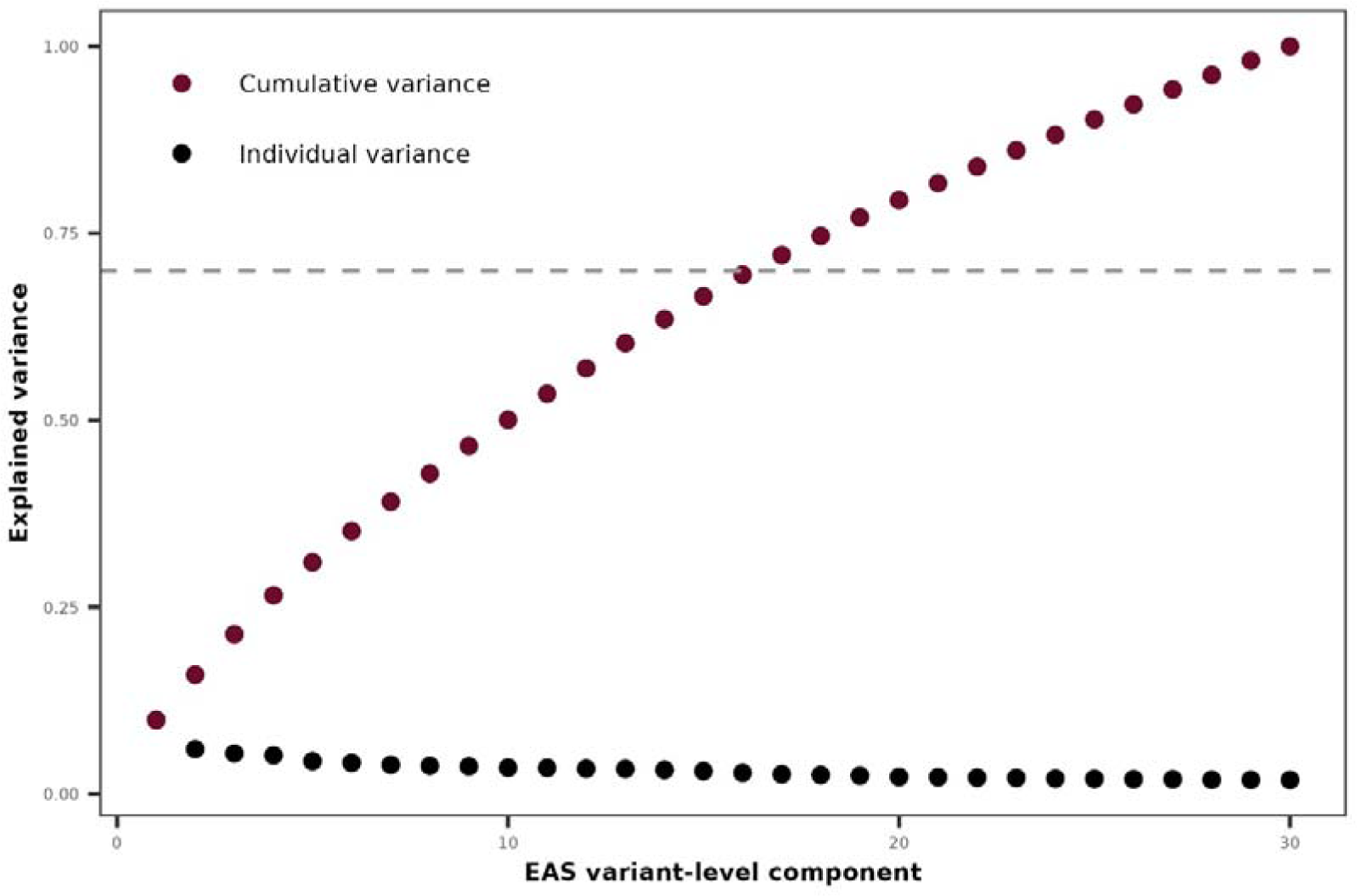
Explained and cumulative variance across latent components. **Note.** Black dots indicate the proportion of variance explained by each latent component, and red dots indicate the cumulative explained variance. The red dashed line marks the 70% cumulative variance threshold, which is reached by the first 16 latent components.

